# Transfer Learning Compensates Limited Data, Batch-Effects, And Technical Heterogeneity In Single-Cell Sequencing

**DOI:** 10.1101/2021.07.23.453486

**Authors:** Youngjun Park, Anne-Christin Hauschild, Dominik Heider

## Abstract

Tremendous advances in next-generation sequencing technology have enabled the accumulation of large amounts of omics data in various research areas over the past decade. However, study limitations due to small sample sizes, especially in rare disease clinical research, technological heterogeneity, and batch effects limit the applicability of traditional statistics and machine learning analysis. Here, we present a meta-learning approach to transfer knowledge from big data and reduce the search space in data with small sample sizes. Few-shot learning algorithms integrate meta-learning to overcome data scarcity and data heterogeneity by transferring molecular pattern recognition models from datasets of unrelated domains. We explore few-shot learning models with large scale public dataset, TCGA (The Cancer Genome Atlas) and GTEx dataset, and demonstrate their potential as meta-learning dataset in other molecular pattern recognition tasks. Our results show that transfer learning is very effective for datasets with a limited sample size. Furthermore, we show that our approach can transfer knowledge across technological heterogeneity, e.g., from bulk cell to single-cell data. Our approach can overcome study size constraints, batch effects, and technological limitations in analyzing single-cell data by leveraging existing bulk-cell sequencing data.

## 1 Introduction

The development of next-generation sequencing (NGS) enables a systematic measurement and analysis of biological questions. During the last decades, a massive amount of biological data is produced. However, at the forefront of the research field, there are still a lack of data issues. The underlying problem arising is that we cannot fully facilitate existing data for new inquiries. There are different reasons for that: (1) each of the datasets is generated under the unique experimental setup, i.e., biological heterogeneity, (2) the batch effect adds noise onto the data in the same experimental setups, and (3) as new sequencing technologies are developed, technical heterogeneity is introduced [1]. Consequently, there is a lack of methodology to integrate various NGS studies. Thus, we need a new methodology to facilitate integrative analysis with pre-existing large-scale biological data to newly produced data for complex biological questions. Thus, the availability of the end-to-end learning with multiple datasets can accelerate this integrative analysis of various data sources [2].

Recently, transfer learning has been introduced to handle biological and biomedical data heterogeneity and integrate different datasets to address novel medical and biological questions. Especially in single-cell omics, these new approaches are used to overcome the technological limitations in this field. Stumpf et al. showed the potential of inter-species knowledge transfer for single-cell sequencing data analysis [3]. BERMUDA introduced autoencoder-based approaches to remove batch effects between datasets in a similar context [4]. Mieth et al. obtained improved clustering results by applying a transfer learning model to a small single-cell sequencing dataset. They transferred knowledge from a large well-annotated source dataset to an unannotated small dataset [5]. In contrast, MARS applied a meta-learning scheme in a single-cell dataset. They trained a deep neural network model with a well-annotated large-scale single-cell sequencing dataset from the TabulaMuris consortium [6] to learn cell type classification ability. In this study the meta-training step improved cell-type classification power and enabled to annotate unknown cell types [7]. Interestingly, the meta-training dataset does not have to contain the same cell types as the test dataset. Meta-training aims to pre-train a model to set initial parameters to learn efficiently from a new, small dataset [8]. This meta-learning approach has already gained attention in cancer genomics, where many large-scale datasets have already been generated compared to other fields [9]. Other studies utilized well-known TCGA, CCLE [10], and functional genomics datasets as a source dataset for meta-training to perform survival analysis [11] and investigate cancer-drug discovery [12].

Although various studies proposed meta-learning and transfer learning approaches to overcome technical limitations in biological and biomedical data, the methodology to overcome technical heterogeneity is not well addressed yet. To fully facilitate our accumulated knowledge in a public database, we must utilize various datasets across different technologies, e.g., bulk-cell sequencing to single-cell sequencing. In this study, we aim to analyse both the potential of meta-learning approaches to overcome data heterogeneity and extremely small sample sizes. Therefore, we train a few-shot learning model to distinguish different expression patterns with only a few examples for unique class labels, i.e., cell or organ types. Meta-training datasets can be large scale NGS datasets about different tissues or organs’ gene-expression profiles, or various case-control datasets. Similar to a few-shot image classification tasks, we expect that the model can recognize unique patterns in a gene-expression profile during meta-training with large-scale and multi-label datasets. Furthermore, we expect to transfer this learned knowledge to different classification tasks from different technologies. Consequently, we show that the model can learn how to handle classification tasks in high-throughput omics biology. This few-shot learning model with a meta-learning algorithm can build a reliable classifier with a marginal amount of data and overcome batch effects and technological heterogeneity in single-cell sequencing projects. In the following, we refer to this few-shot classification model as a transferable molecular patterns recognition model (TMP).

## 2 Results

### 2.1 Meta-learning for recognizing transferable molecular patterns and handling of small-size data

In this study, we used a two-step learning phase, namely the meta-training and the fine-tuning phase (Figure 1a). In the meta-training phase, the model learns how to recognize gene expression patterns from large-scale transcriptome data. The data for fine-tuning does not have to be in a similar context or the same technology compared to the data used for meta-training. However, it needs to be within a similar context or similar sequencing technology for adjusting batch effects or technological heterogeneity.

**Figure 1:**
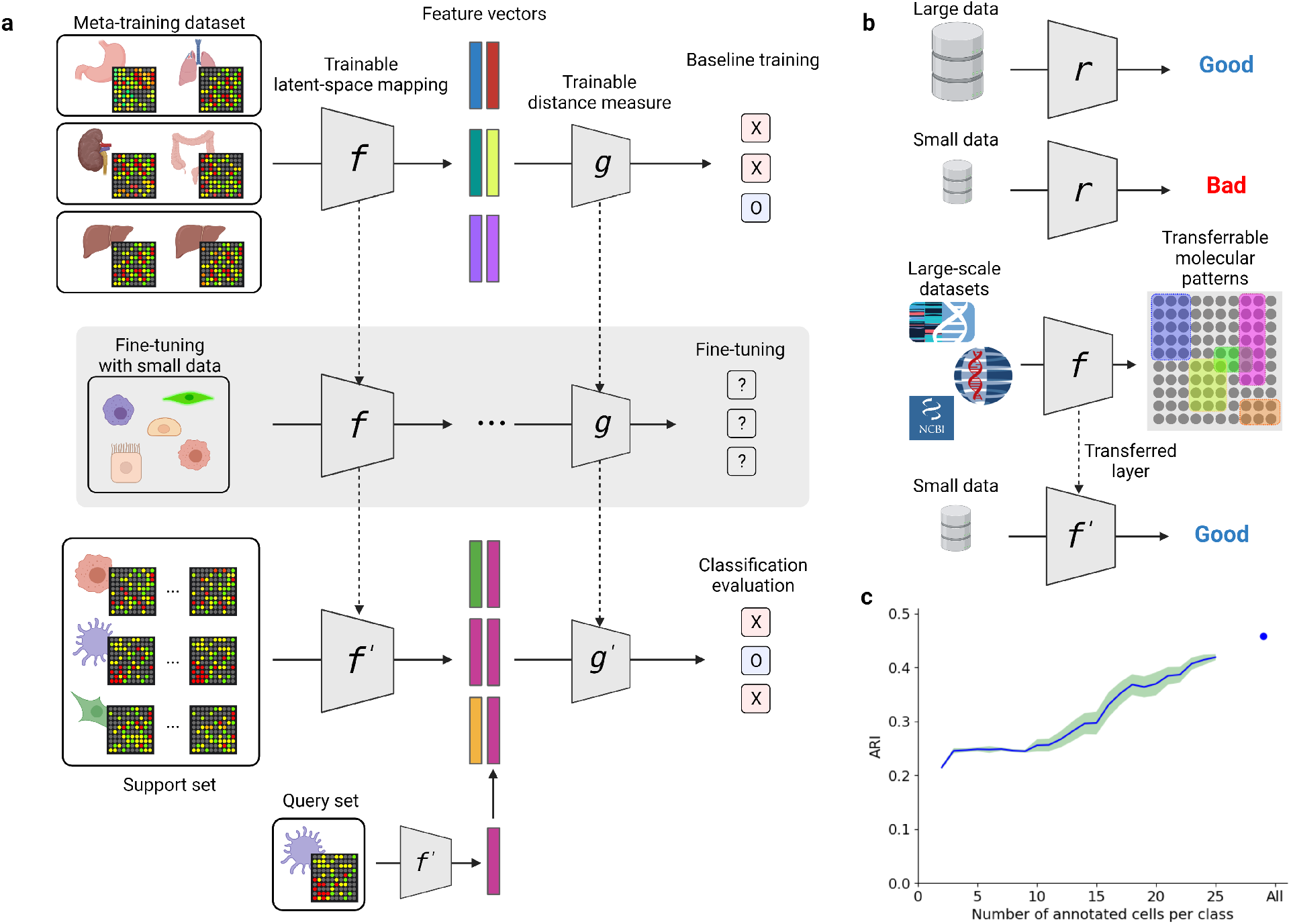
Overview of our transfer learning scheme. **a.** Training and testing schemes of our meta-learning approach. **b.** Hypothesis of this study. Transfer learning with large-scale datasets can improve models based on small datasets and different sequencing technologies. **c.** Adjusted rand indexes (ARI) for PCA-based clustering in cell type classification problems in single-cell sequencing datasets. The blue line is an average score, and the light green area is 95% confidence interval.

The machine learning model requires a reasonable amount of data for the training phase to ensure a good performance in terms of accuracy (Figure 1b). In the single-cell sequencing data field, many approaches use dimensionality reduction methods to overcome the technical limitation of single-cell sequencing itself. For example, recent studies employed PCA (Principal Component Analysis), tSNE (t-distributed stochastic neighbor embedding), or UMAP (Uniform Manifold Approximation and Projection). Other recent methods are based on autoencoders and intend to reduce noise and batch effects and embed single-cell gene expression profiles into feature vectors with reduced size. Those approaches can be very effective when the given dataset has well-balanced and large enough samples. In our study, we used PCA and k-means clustering with single-cell data to evaluate the impact of data size on a model. We aimed at keeping this analysis as simple as possible and thus we did not use transductive inferences such as tSNE or UMAP. We used a randomly picked small dataset to calculate the principal components based on human pancreas single-cell data. After that, all samples are projected to the latent space and clustered subsequently (details in the Methods). Until data size reaches ten samples per class, the ARI (Adjusted rand indexes) score did not change. When more data is used for finding the latent space, the k-means clustering results gradually improved (Figure 1c). In the following, we will focus on the ‘small-size data’ condition and show that meta-learning and fine-tuning training can improve a model having limited data, which is a crucial limiting factor in many areas of biomedical research.

### 2.2 Overview of our TMP model

TMP is based on a relation network model for few-shot image classification [13] which comprises two trainable deep neural networks (DNN). The first network projects the given gene expression profile into the feature vector, and the second network measures the distance between two given feature vectors. Both DNNs are trained by backpropagation to find proper latent spaces and relevant distance measures for the latent spaces during the meta-training step. Given large-scale multi-class gene expression profile datasets, TMP learns how to embed given gene expression data and measure distances between data points in latent space. Afterwards, this pre-trained TMP is fine-tuned with a small amount of data from the target task (Figure 1a). Details about the network can be found in the Methods section.

In this study, we pre-trained the model with GTEx data, i.e., human tissue-level gene expression profiles. Based on this pre-trained model, we adapted it for two different datasets: (1) TCGA cancer dataset to analyze the few-shot learning model concerning its ability to recognize transferable molecular patterns, and (2) to human single-cell pancreas datasets to show the possibility of cross-modal data integrative analysis. TMP can learn gene expression patterns from GTEx, thus can be directly applied for other datasets and tasks. Furthermore, with these adjusted parameters, the model can quickly be adapted to a new task. Our approach demonstrates that batch effects or technical heterogeneity in single-cell omics can be handled and compensated with TMP.

### 2.3 Transfer learning can improve classifiers built with small-size data

Our few-shot learning model aims to solve a variety of classification problems with limited data. We transferred knowledge from a large-scale dataset that is not directly related to the classification problem to achieve the goal. To investigate the general performance of the few-shot learning model on the transcriptome dataset, we used TCGA and GTEx datasets which are large-scale omics with multi-class labels datasets. TCGA has a human organ-level class label, and GTEx has a human tissue-level class label. Thus, the two datasets are in a similar context but have different class label sets, which makes them ideal candidates for few-shot learning [14]. First, we trained our model with the GTEx dataset. With this meta-training step, the model learns to classify different types of tissues by gene expression patterns. Without any additional fine-tuning, the GTEx meta-trained model shows 78.91 % ± 0.76 % accuracy on TCGA. The TCGA meta-trained model shows 84.57 % ± 0.66 % accuracy on the GTEx data in a 5-way 5-shot task (Table S3. Next, we compared the cancer type classification problem with recently published methods. We fine-tuned the model with only 15 samples per class and tested with the remaining (495 samples out of 9,781; 5% of the dataset). Although the accuracy of the model can be affected by the randomly chosen 15 samples, we could obtain an average accuracy of 94%+ in 10 repeated training runs, even though we used only 5% of the TCGA data for fine-tuning (Figure 2a). Although accuracy in few-shot learning is not directly comparable to standard models, our approach shows similar performance (95.6% and 95.7% accuracy) compared to recent CNN-based classification models that used 80% of the data for training [15].

**Figure 2:**
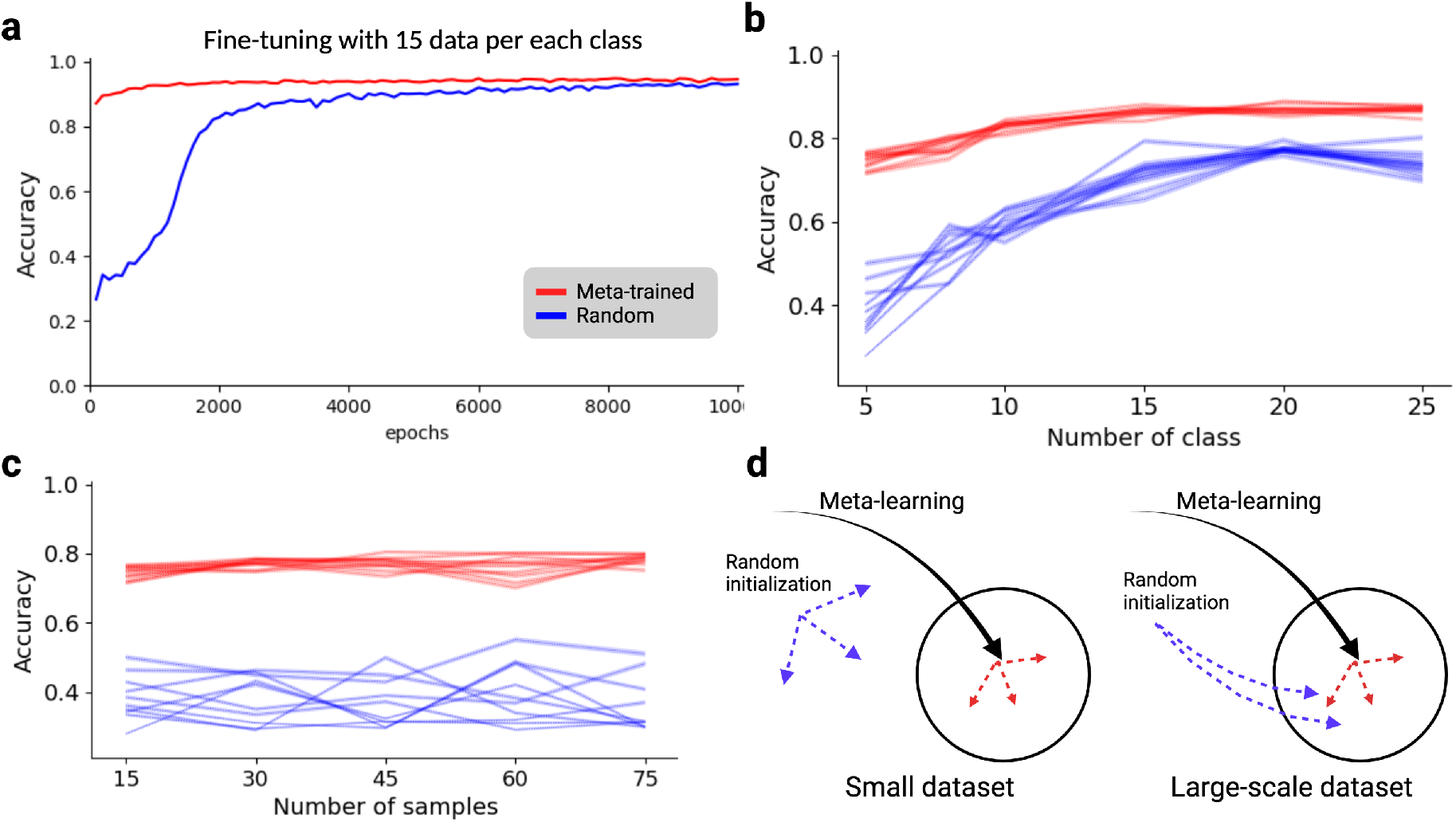
An accuracy chart of 5way-5shot task with TCGA dataset. **a.** An 5-way 5-shot accuracy chart during fine-tuning with TCGA pan-cancer dataset. Both red and blue lines used 15 samples for each of 33 different cancer types. The red line is meta-trained with GTEx dataset, and the blue line is randomly initialized. **b.** Accuracy score varying number of classes and samples after fine-tuning. Red lines are meta-trained models and blue lines are randomly initialized models. Dotted line is average score of tests and colored are is a 95% confidence interval. **c.** Accuracy score varying number of classes and samples after fine-tuning. Red lines are meta-trained models and blue lines are randomly initialized models. Number of samples is fixed to five. Dotted line is average score of tests and colored are is a 95% confidence interval. **d.** Illustration to compare learning processes for both variables, meta-training and given data sizes in fine-tuning.

When using all 33 classes in the TCGA cancer dataset, the meta-trained model converged early in comparison to the randomly initialized model which needs more epochs to converge. However, fully-trained both models have similar accuracy (Figure 2a). With this fully supervised setup, we can see that transferred knowledge can reduce search space and help to reach early convergence. To investigate transferred knowledge as general information that can enhance the few-shot learning model, we trained the model with a subset of the samples of the 33 classes and tested with all remaining samples. We compared two baseline models: meta-trained with GTEx (1) and a randomly initialized model (2) and further trained them with 5, 10, 15, 20, and 25 classes out of the 33 classes. This fine-tuning is stopped when batch accuracy reached 0.99% to avoid overfitting (Methods). With this result, we can observe that transferred knowledge is very useful when fine-tuning data is limited to a small number of different classes. With a meta-trained model, we can build a few-shot model with 80%+ accuracy with only 1.5% of the TCGA dataset (10 out of 33 classes and 15 samples per class). Because the model agnostic meta-learning (MAML) algorithm is efficient with small data sizes, a more critical factor in better accuracy was the number of classes than the number of samples. When we fixed the number of classes to five and varied the number of samples from 15 to 75, the accuracy change was not significant (Figure 2c). With the ARI measure, it shows similar patterns (Figure S1). However, if the number of classes is 25, we can see marginal improvement when the number of samples is increased (Figure S2). The analysis of the TCGA and GTEx datasets, demonstrates that few-shot learning with a meta-learning algorithm can be a new method to facilitate existing large-scale datasets and transfer knowledge. Model initialization with transferred knowledge can improve model accuracy and also reduce training times (Figure 2d).

### 2.4 Meta-learning across technical heterogeneity

TCGA and GTEx can not fully reflect the current technological heterogeneity issue in omics biology. Thus, we further investigated a single-cell sequencing transcriptome dataset. Various methods have been introduced for effective batch correction and integrative analysis with multiple datasets from various techniques in this field. In our study, we used five different single-cell datasets from the human pancreas and the GTEx and TCGA datasets as meta-training datasets. Pretrained model with both large-scale datasets shows accuracies of less than 50% accuracy on single-cell sequencing datasets (Table S3). Based on this test results from few-shot baseline models with GTEx and TCGA datasets, we could show that the models can identify different cell types from single-cell sequencing datasets even though a model has been trained only as a organ/tissue level classification task. However, test classification result shows that the meta-trained models recognize same class label in different dataset to different class label (Figure S4).

Next, we used the Baron dataset out of five different single-cell datasets for fine-tuning of the GTEx-pretrained model to validate that our model can transfer knowledge from a large-scale bulk-cell sequencing dataset to a single-cell problem. We obtained accuracy from five different training conditions (Figure 3a): meta-trained and fine-tuned with the Baron dataset (filled circle), randomly-initialized and trained with the Baron dataset (hollow circle), meta-trained and fine-tuned with subset of the Baron dataset (filled square), randomly-initialized and trained with subset of the Baron dataset (hollow square), and meta-trained (filled diamond). Similar to the results from the TCGA data, meta-training does not significantly affect the accuracy if a fine-tuning dataset has enough data. However, if the fine-tuning data is limited, meta-training can improve the quality of the model significantly. During training with Baron data, we observed that the accuracy converges within 100,000 epochs (Figure 3b). In these single-cell datasets, the classification accuracy in other similar single-cell sequencing datasets is improved if a fine-tuning training uses more than 15 samples from Baron (Table S4). This improvement of accuracy was directly observable in the detailed result of test episodes. After learning the cell types classification problem with the baron dataset, the few-shot model was able to distinguish different cell types more accurately. Furthermore, we can identify groups of similar cell types across different datasets (Figure S5). Notably, we could observe interesting classification results in the heatmap such as ‘alpha-contaminated’ and ‘beta-contaminated’ in the Xin dataset, or ‘dropped’ in Wang dataset, ‘co-expression cell’ and ‘unclassified endocrine cell’ in the Segerstolpe dataset. Because few-shot model can compare query sample to more class labels at one time in a 20-way and 5-shot task, we can see distinct clusters of similar labels and cell clusters that are not clearly annotated.

**Figure 3:**
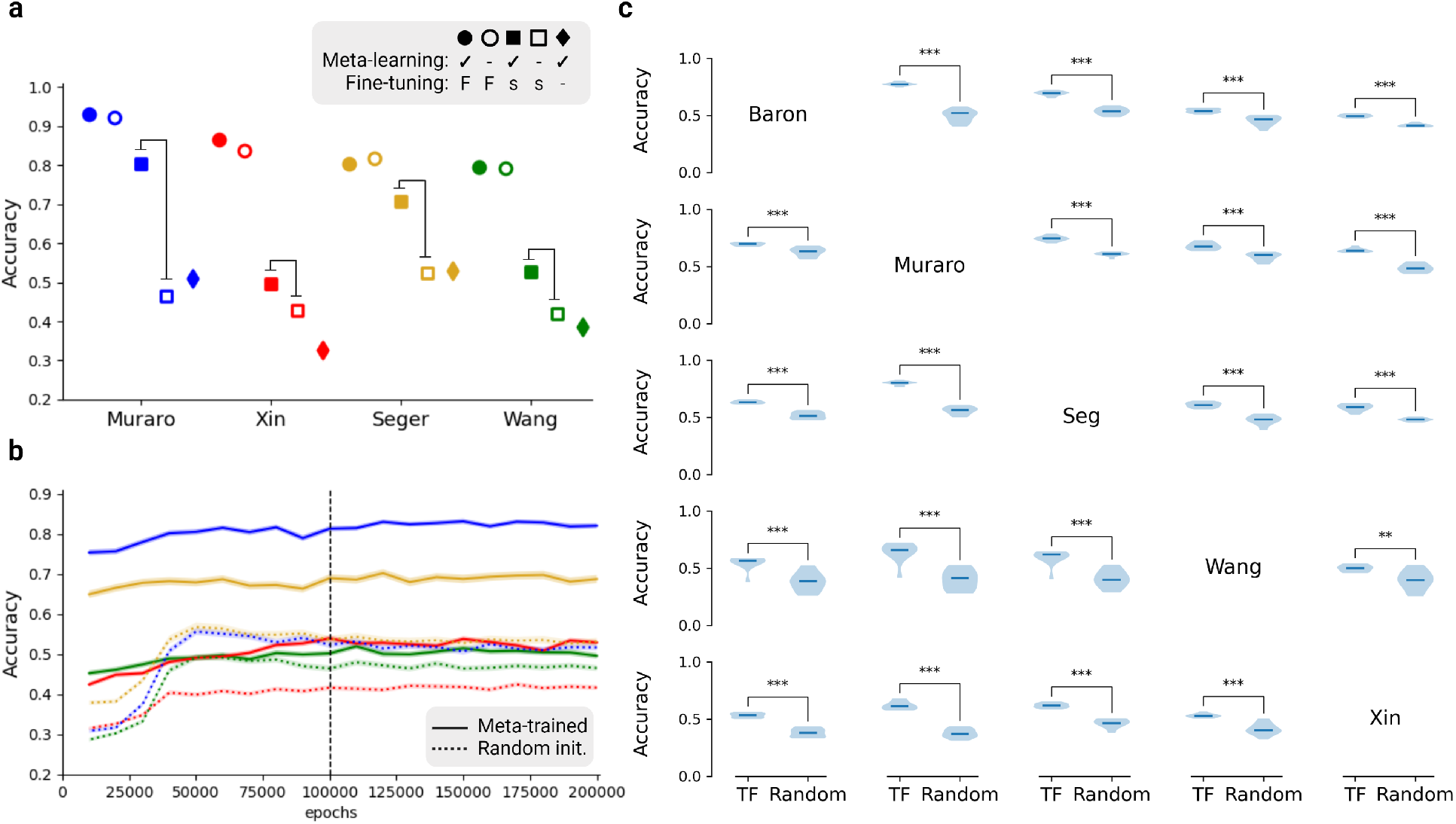
An accuracy chart of 5way-5shot task with five different single-cell pancreas datasets. **a.** Comparisons between different training conditions show that transfer learning makes a significant difference in small-size data conditions. Meta-learning refers to pre-training with the GTEx dataset, and it is represented with filled markers. Fine-tuning refers to additional training with the Baron dataset. ‘F’ indicates that the complete dataset of Baron was used for the fine-tuning training step, and it is represented with a circle. ‘s’ indicates a data scarcity condition, and it is represented with a square. ‘-’ indicates no fine-tuning training, and it is represented with a diamond. (b) and (c) figures are about the data scarcity condition (filled and hollow square cases). **b.** Accuracy chart during training for fine-tuning with subset of the Baron dataset (15 samples per class) in case for two squares in (a). After 50,000 epochs, performance of the model converged. Solid lines represent models based on meta-trained networks with GTEx dataset and further trained with a subset of the Baron dataset. Dotted lines are randomly initialized networks trained with 15 samples per class from the Baron dataset. **c.** Pair-wise comparisons of 5 different single-cell pancreas datasets show accuracy differences between meta-trained and randomly initialized networks. Each of the violin plots is composed of 10 repeats of training. ‘TF’ indicates fine-tuning started with meta-trained network and ‘Random’ indicates randomly initialized models. The rows represent the dataset used for fine-tuning, the column the dataset used for testing. Statistical significance evaluated by Student’s t-test is indicated by *: *p* < 0.05, **: *p* < 0.01, ***: *p* < 0.001.

We further investigated the fine-tuning with small-size data, here 15 data per class (see Figure 3a). Thus, we compared all possible scenarios with single-cell datasets. We chose one dataset out of five as a fine-tuning dataset and tested its 5-way 5-shot task accuracy with the others. Meta-trained models and randomly initialized models are fine-tuned with 15 samples per class from the chosen dataset. Because the data quality of randomly picked 15 cells can significantly affect the training, we repeated the training 10 times and tested for significant differences. It turned out that the meta-trained model is significantly better than the standard model when the available training data is limited (Figure 2c).

## 3 Discussion

As next-generation sequencing is routinized in experimental biology, the speed of data accumulation has been accelerated. However, most studies can utilize only partial data because the biological heterogeneity, batch effects, and technical heterogeneity, which hinder integrative analysis. Thus, every study has to produce large amounts of data to obtain a significant result. Few-shot learning with meta-learning algorithms could solve this issue in biomedical research and has already been used in other areas, e.g., computer vision [14]. Thus, we propose that a few-shot learning model with a meta-learning algorithm can be used as a new methodology to exploit existing public resources and data to build reliable models for small sample size. This approach is not only cost but also time-efficient, and thus, very important for biomedical studies, e.g., for cancer. In a recent cancer-drug discovery study [12], meta-learning has been used for functional genomics data to extract biological knowledge from very small sizes of screening data. Consequently, by using this new approach, we can investigate biological systems in higher resolution and lower costs [9].

In this work, we used a relatively simple structure of networks for the omics data compared to applications in computer vision in order to demonstrate that transfer learning is a reliable method for handling of biological heterogeneity, batch effects, and technical heterogeneity in omics analyses. However, there is room for improving the models for various data integration analyses. We expect that a more complex but well-optimized structure for biological data can improve knowledge transfer and expand the applications also to other sequencing variations. Here, we analyzed single-cell sequencing datasets as an example of a technological heterogeneity issue. There are already many successful models for the integration of single-cell sequencing datasets; Seurat [16], scVI [17], scVAE [18], Scanorama [19], MARS [7], and scETM [20]. Those methods, however, focused on only single-cell data integration. Our findings indicate that besides single-cell datasets, other types of data e.g., existing bulk-cell datasets, are useful for interpreting single-cell sequencing datasets. Moreover, we expect that our cross-technology transfer learning scheme can also be applied to those recent state-of-the-art batch correction models. Based on our findings, we are hypothesise that meta-learning on large bio-molecular datasets in combination with fine-tuning can improve model performance in many applications struggling with small sample sizes.

## 4 Methods

### 4.1 Data preprocessing

We downloaded processed data of TCGA, GTEx, and five single-cell datasets (section in Data Availability). We removed the ambiguous label, ‘not applicable’, in the Segerstolpe dataset. Depending on its technical origin, each dataset has a different set of genes. Thus, we need to fix the gene list for further analysis. 18,000 genes are selected from the intersection of TCGA, GTEx, and StringDB. Because our goal is not mining novel genes for cell-type annotation, StringDB is applied as the simplest feature selection method to control memory usage and training speed. Different gene-IDs are converted to gene symbols with the Biomart package, and missing genes in the independent dataset are filled with 0. After filtering the gene list, we Log2-normalized the gene expression levels and re-scaled it to a 0-1 scale.

### 4.2 Implementation few-shot

#### Transferable molecular pattern recognition model network structure

Our network model has two trainable networks, the feature encoder and the relation network [13]. Feature encoder blocks are composed of a fully connected layer, a batch normalization layer, and a ReLU layer. Feature encoder for a gene expression profile of 18,000 genes is composed of three blocks of feature encoders. Additionally, the feature encoder is a non-negative network by clamping negative weight to zero during iterations to focus on co-expressional patterns rather than other regulatory mechanisms. This approach is inspired by work with non-negative kernel autoencoders [21]. The relation network block is composed of three blocks. Two blocks are composed of a fully connected layer, a batch normalization layer, and a ReLU layer. The last block is composed of a fully connected layer and a sigmoid function for the output value.

The feature encoder (*f*(*x*)) has three encoding blocks. Each of the fully connected layers is set to 4000, 2000, and 1000 neurons. The relation network (*r*(*x*)) is set to 500 and 100 neurons. Consequently, the 18,000 genes are embedded into 1,000 length size vectors through the feature encoder, and the relation network gets two feature vectors for comparison.

In training and evaluation, we used the same design with a support set 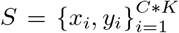 and a test (or query) set 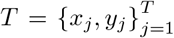 from given dataset. We randomly selected five classes (C=5) for each training iteration and five samples for each class (K=5) for a support set. In the same class set, we randomly selected ten samples (T=10; training and testing batch sizes) for a training/testing batch set. With these datasets, loss is defined as follows:

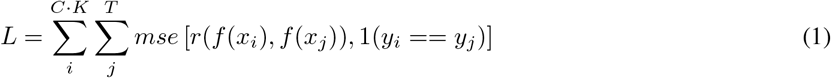

After each iteration, we clamped negative weights in the feature encoder to zero. We use the Adam optimizer with a learning rate of 0.0005, and every 100,000 epochs decreased the learning rate by half during a meta-training phase. We stopped pre-training the model at 300,000 epochs and transferred this network onto different datasets. For fine-tuning, we used the Adam optimizer with a learning rate of 0.0001. The results in Figure 3c reflect the evaluation scores at 100,000 epochs of fine-tuning. When we use larger batch sizes (T) and a greater learning rate, fine-tuning can stop at less than 100 epochs.

#### Few-shot training with GTEx

The GTEx dataset was used as a meta-training dataset in both TCGA data analysis and single-cell pancreas datasets analysis. To ensure that meta-training has been carried out with sufficient samples, we held out tissue with less than 100 samples, i.e., Bladder (21), Cervix Ectocervix (9), Cervix Endocervix (10), Fallopian Tube (9), Kidney Medulla (4), and Kidney Cortex (85). We randomly selected five tissues during the training phase and selected five examples as support set and ten samples as training batch query set.

#### Few-shot training with TCGA

To investigate a few-shot learning task in omics data, we used a meta-trained model with the GTEx dataset and fine-tuned it with the subset of TCGA data. The fine-tuning dataset (15 samples per class) was randomly selected and tested with the others. The accuracy is obtained from 5,000 episodes with 10 test batch sizes (Figure 2a). To further investigate the data quantity issue, we evaluated various classes and numbers of data out of 33 cancer types and their samples. Because some of the classes in the TCGA dataset has less than 75 samples, we excluded the cancer types with less than 100 samples in the training step. Training with the TCGA dataset was done until 50,000 epochs with step size 1,500. In every 2,500 epochs, we checked batch accuracy and early stopped the training. If the average batch accuracy of 2,500 epochs is greater than 0.99, the fine-tuning was stopped. Evaluation is done with 1,000 episodes with 10 test batch sizes. We did ten repeats with random seed numbers from 1 to 10.

#### Few-shot training with single-cell pancreas data

For the batch effect and technical heterogeneity task, five datasets were split such that one dataset was used for training, and the remaining datasets were held for evaluation. We removed cells which have not enough samples for training and testing in a 5-way 5-shot task, namely t_cell (7), schwann (13), epsilon (18) in Baron, epsilon (3), unclear (4) in Muraro, MHC class II cell (5), mast cell (7), epsilon cell (7), unclassified cell (2) in Segerstolpe, acinar (6), delta (9) in Wang, PP.contaminated(8) and delta.contaminated(9) in Xin. In fine-tuning, we selected 15 samples per class in the training datasets. The evaluation has been carried out by 5,000 episodes with 10 test batch sizes. Average and 95% confidence interval are represented in (Figure 3b). Raw values are available in the GitHub repository. Each sample was randomly selected, thus we repeated this training ten times and reported the distribution of the average value over these replicates (Figure 3c). Additionally, the average value and 95% confidence interval for each run are available in the GitHub repository.

### 4.3 Conventional machine learning methods

We used the Python package scikit-learn http://scikit-learn.org/stable/index.html to implement conventional machine learning methods.

For PCA and K-means clustering, we randomly selected five classes and a small subset of data per class from 2 to 25 for training. All other data points are projected on the latent space, and we did k-means clustering to find five clusters. Because K-means is starting with random initialization, we did 100 repeats for each data point. The performance is evaluated with an adjusted rand index (ARI), and average ARI with 95% confidence intervals are shown.

## 5 Data Availability

The datasets generated during and/or analyzed during this study are all public data: TCGA (https://portal.gdc.cancer.gov/); GTEx (https://gtexportal.org/home/index.html); Size of TCGA dataset is summarized in Table S1. Single-cell pancreas datasets are summarized in Table S2. It can be downloaded at Hemberg-lab’s github (https://github.com/hemberg-lab/scRNA.seq.datasets).

## 6 Code availability

All the code used in this study is written in Python using scikit-learn and PyTorch library. The source code, figures, and code for figures are available on Github at https://github.com/iron-lion/tmp_model.

## Acknowledgments

This project has received funding from the European Union’s Horizon2020 research and innovation programme under the grant agreement No 826078. This publication reflects only the authors’ view and the European Commission is not responsible for any use that may be made of the information it contains.

## 7 Supplementary Figures

S1 ARIs of training results with TCGA datasets

S2 Fine-tuning training results using 25 classes of TCGA datasets

S3 ARI of pair-wise comparisons of 5 different single-cell pancreas datasets

S4 Heatmap of test results of the meta-trained model in single-cell datasets

S5 Heatmap of test results after the fine-tuning

**Figure S1:**
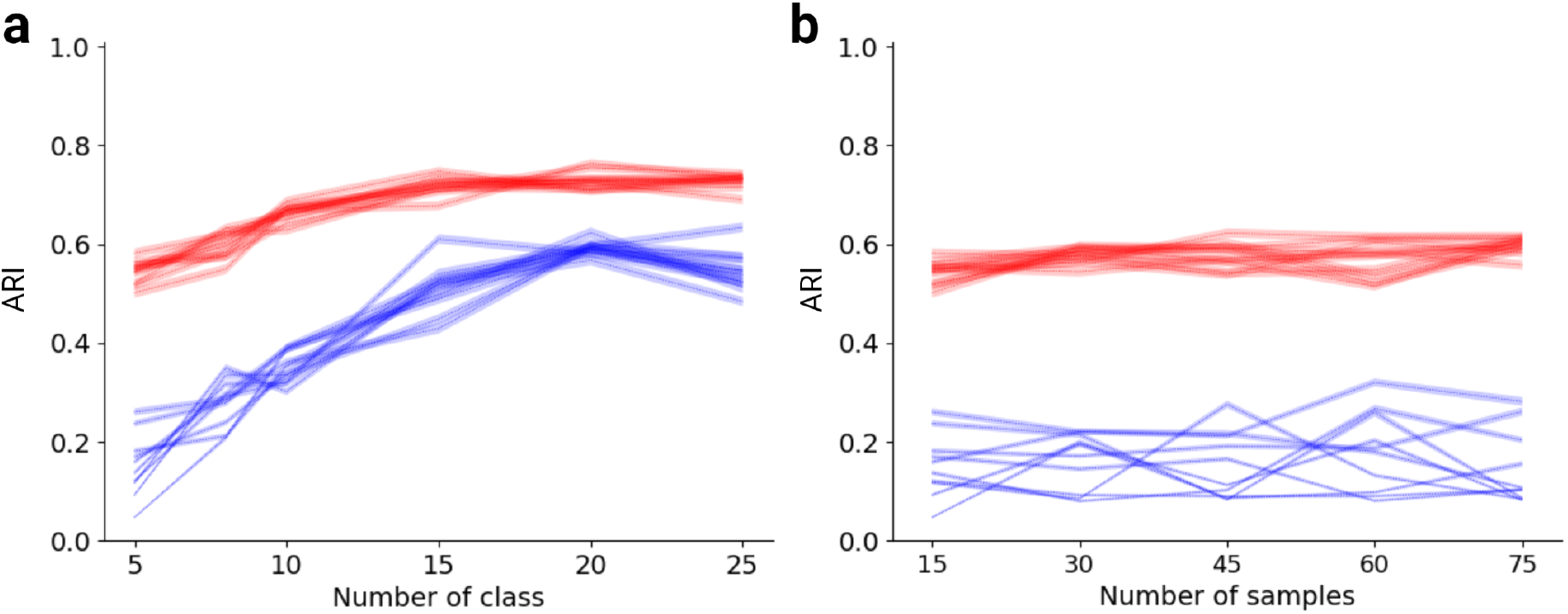
ARIs of training results with TCGA datasets. **a, b.** ARI score of (Figure 2b,c). In (a), the number of samples is fixed to 15. In (b), the number of the class is fixed to five. Training is done with baron data. Because training results are influenced by set of samples randomly picked, this results are obtained from ten repeats. The dotted line is average ARI of test episodes for each of ten repeats and the colored area represents 95% confidence interval.

**Figure S2:**
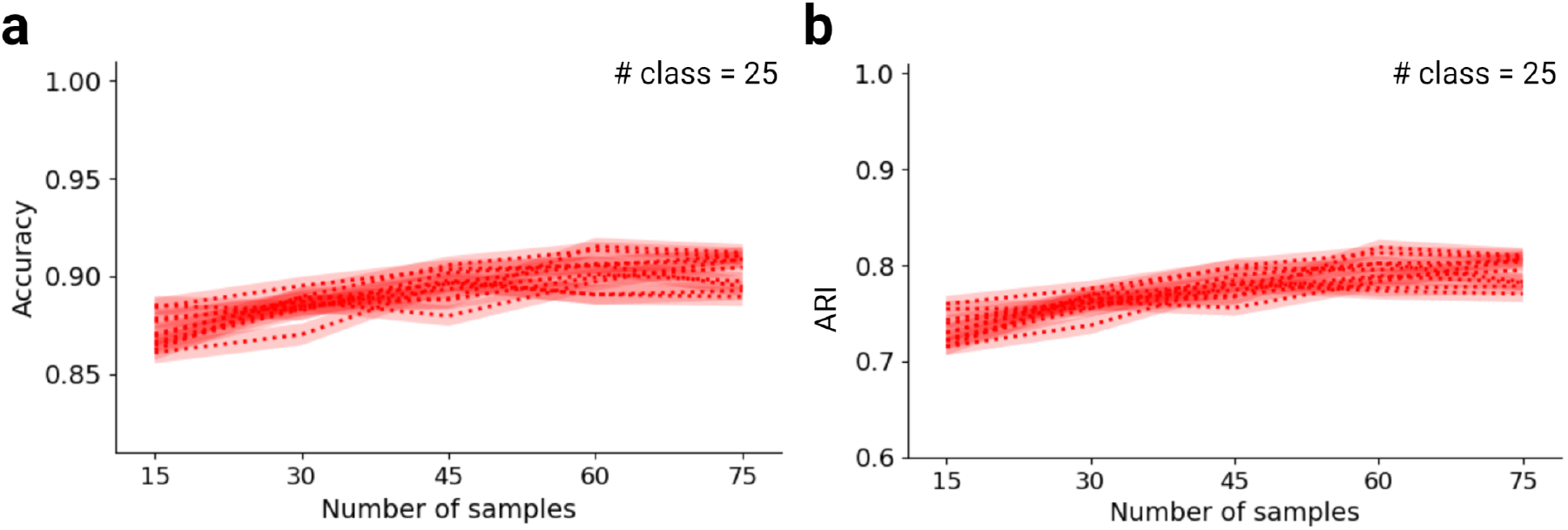
Fine-tuning training results using 25 classes of TCGA datasets. **c, d.** The model is pre-trained with the GTEx dataset, and fine-tuning is done with a fixed number of classes to 25. The accuracy and ARI marginally improved the following number of samples. Ten repeats of the test are done with test episode 1,000 and test batch sizes ten. The dotted line is average accuracy (c) or ARI (a, b, d) of test episodes for each of ten repeats and the colored area represents 95% confidence interval.

**Figure S3:**
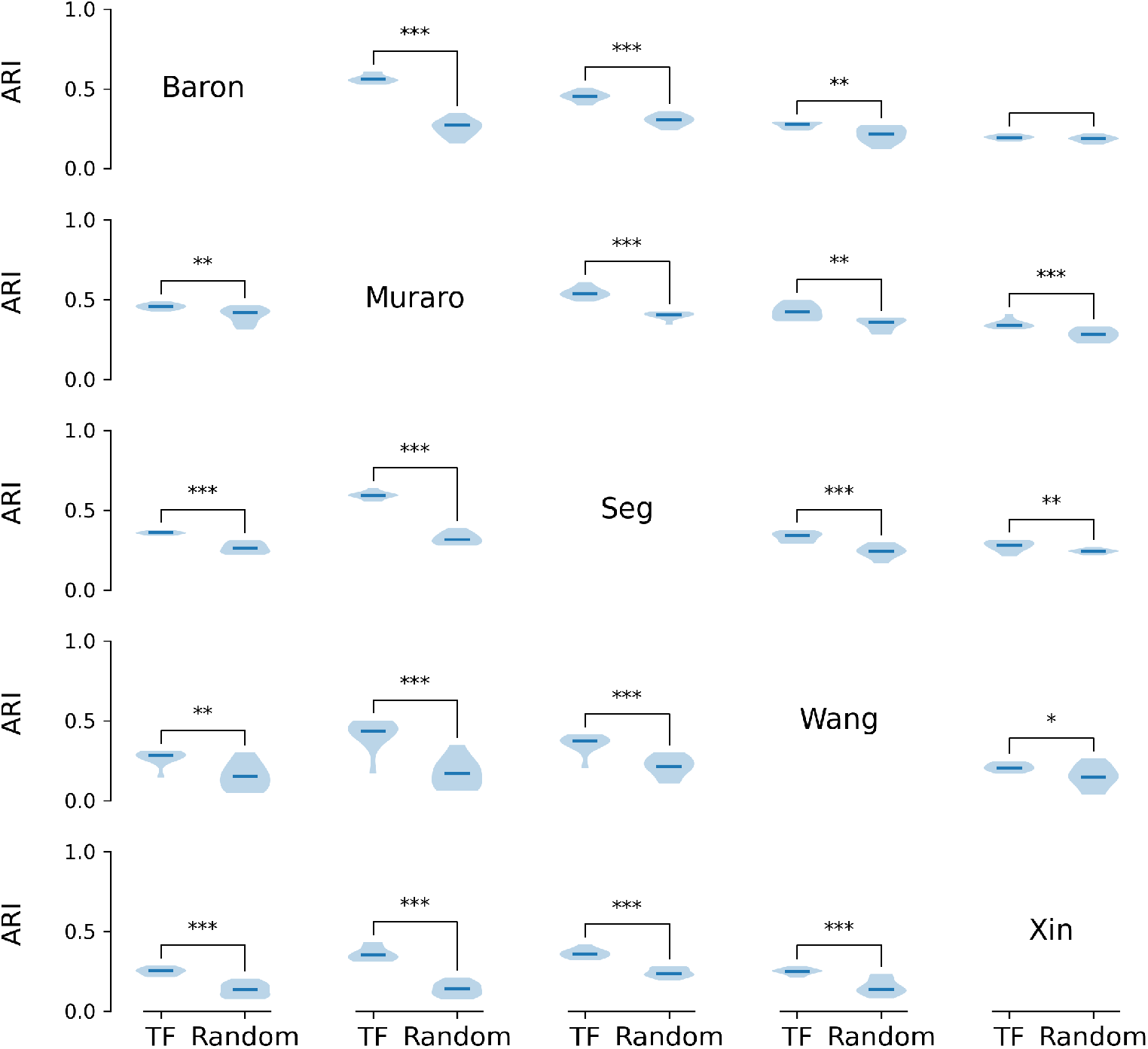
ARI of pair-wise comparisons of 5 different single-cell pancreas datasets. Pair-wise comparisons of 5 different single-cell pancreas datasets show ARI differences between meta-trained and randomly initialized networks when they further trained with 15 samples per class in each of data. The rows represent the dataset used for fine-tuning, the column the dataset used for testing. Each of the violin plots is composed of 10 repeats of training. ‘TF’ indicates fine-tuning started with the meta-trained network and ‘Random’ indicates randomly initialized models. Test batch size was ten in a 5-way 5-shot task. Thus, at each test episode, this model classifies 50 samples and evaluates the ARI score. Statistical significance evaluated by Student’s t-test is indicated by *: *p* < 0.05, **: *p* < 0.01, ***: *p* < 0.001.

**Figure S4:**
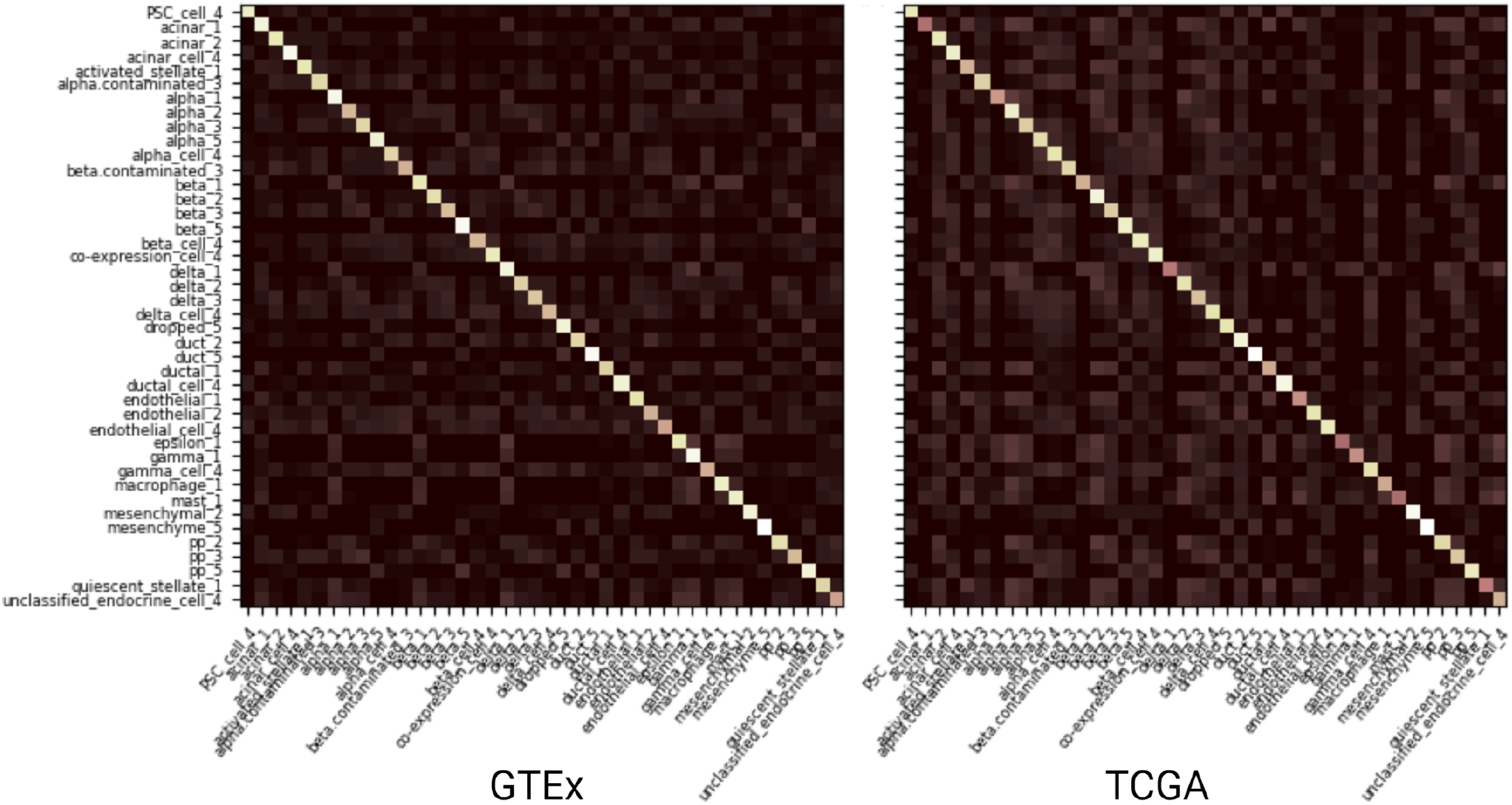
Heatmap of test results of the meta-trained model in single-cell datasets. We merged all five datasets into one test set, and did 10-way 5-shot test with 10,000 episodes. Row index is true label and column index is predicted label. The suffix number indicates the different datasets (1: Baron, 2: Muraro, 3: Xin, 4: Segerstolpe, 5: Wang).

**Figure S5:**
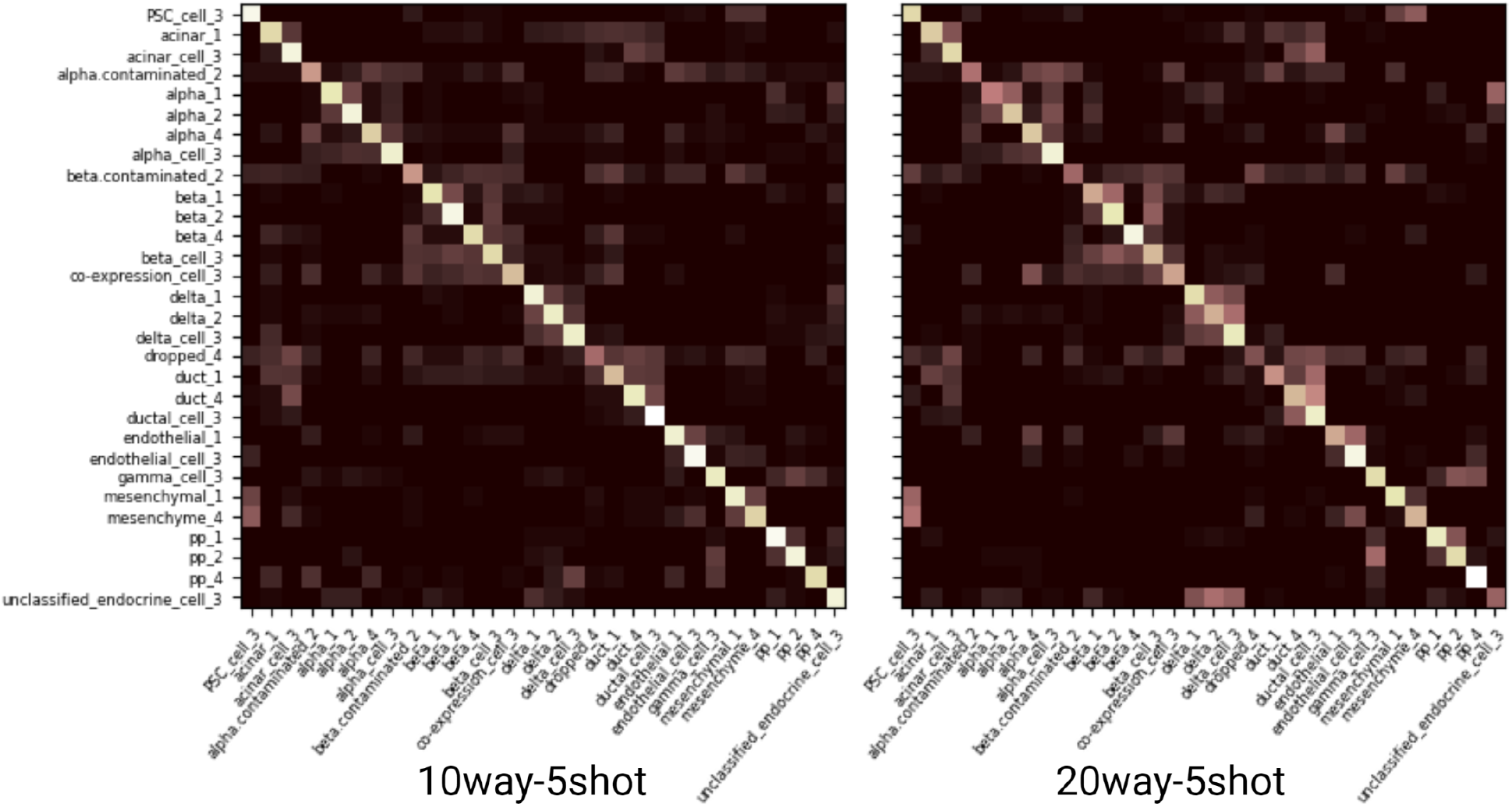
Heatmap of test results after the fine-tuning. We used a meta-trained model with the GTEx dataset and further trained with the Baron dataset. We merged the remaining four datasets into one test set and did 10-way 5-shot and 20-way 5-shot tests with 10,000 episodes. In the 20-way 5-shot task, the clusters of similar cell types in different datasets are more clearer. Row index is true to label and column index is predicted label. The suffix number indicates the different datasets (1: Muraro, 2: Xin, 3: Segerstolpe, 4: Wang).

## 8 Supplementary Tables

S1 Downloaded TCGA cancer dataset description.

S2 Single-cell human pancreas data description

S3 Baseline accuracy for GTEx and TCGA pre-trained model

S4 Accuracy for the single-cell human pancreas data

**Table S1:** Downloaded TCGA cancer dataset description. We used all 33 cancer types in TCGA. The number includes non-tumor samples and we exclude normal samples for further analysis.

| Cancer type | Files | Cancer type | Files | Cancer type | Files | Cancer type | Files |
| --- | --- | --- | --- | --- | --- | --- | --- |
| PAAD | 177 | STAD | 375 | SARC | 262 | LAML | 151 |
| KICH | 65 | UCEC | 552 | UVM | 80 | THYM | 119 |
| PRAD | 498 | COAD | 479 | TGCT | 150 | LUAD | 535 |
| UCS | 56 | DLBC | 48 | LUSC | 502 | THCA | 502 |
| BRCA | 1102 | LIHC | 374 | LGG | 529 | PCPG | 178 |
| OV | 379 | ESCA | 161 | ACC | 79 | CHOL | 36 |
| BLCA | 414 | KIRC | 538 | KIRP | 288 | SKCM | 103 |
| GBM | 169 | READ | 167 | CESC | 304 | MESO | 86 |
| HNSC | 500 |  |  |  |  |  |  |

**Table S2:** Single-cell human pancreas data description.

| Dataset | Size | Class | Protocol | Accession ID |
| --- | --- | --- | --- | --- |
| Baron (Human) [22] | 8,569 | 14 | inDrop | GSE84133 |
| Muraro [23] | 2,126 | 10 | CEL-Seq2 | GSE85241 |
| Segerstolpe [24] | 2,209 | 14 | Smart-Seq2 | E-MTAB-5061 |
| Xin [25] | 1,600 | 8 | SMARTer | GSE81608 |
| Wang [26] | 457 | 7 | SMARTer | GSE83139 |

**Table S3:** Baseline accuracy for GTEx and TCGA pre-trained model. Each of the transferred models is trained with both GTEx and TCGA datasets. Accuracy is obtained from 1,000 test episodes. A 95% confidence interval of 1,000 test episodes is shown.

| Transfer | Accuracy | Transfer | Accuracy |
| --- | --- | --- | --- |
| GTEx -> TCGA | 78.91 % ± 0.76 % | TCGA -> GTEx | 84.58 % ± 0.66 % |
| GTEx -> Baron | 43.69 % ± 0.54 % | TCGA -> Baron | 45.50 % ± 0.53 % |
| GTEx -> Muraro | 50.87 % ± 0.57 % | TCGA -> Muraro | 45.01 % ± 0.53 % |
| GTEx -> Xin | 32.30 % ± 0.40 % | TCGA -> Xin | 32.75 % ± 0.41 % |
| GTEx -> Segerstolpe | 52.88 % ± 0.63 % | TCGA -> Segerstolpe | 49.44 % ± 0.57 % |
| GTEx -> Wang | 38.71 % ± 0.48 % | TCGA -> Wang | 37.30 % ± 0.47 % |

**Table S4:** Accuracy for the single-cell human pancreas data. The model was pre-trained with the GTEx dataset and fine-tuned with Baron data. Each model was trained with 15, 30, and 45 samples per class and the entire Baron dataset. We evaluated the model after fine-tuning 100,000 epoch with a learning rate of 0.0001. Accuracy and ARI are measured from 1,000 test episodes in the 5-way 5-shot classification task.

| Test dataset |  |  | 15 | 30 | 45 | 100% |
| --- | --- | --- | --- | --- | --- | --- |
| Muraro | Random | Accuracy | 35.6 % ± 0.5 % | 48.4 % ± 0.6 % | 56.9 % ± 0.5 % | 91.2 % ± 0.4 % |
| Muraro | TF | Accuracy | 81.4 % ± 0.5 % | 83.6 % ± 0.5 % | 86.3 % ± 0.4 % | 92.3 % ± 0.4 % |
| Muraro | Random | ARI | 0.090 ± 0.003 | 0.164 ± 0.004 | 0.270 ± 0.005 | 0.806 ± 0.007 |
| Muraro | TF | ARI | 0.628 ± 0.008 | 0.666 ± 0.008 | 0.706 ± 0.007 | 0.835 ± 0.007 |
| Xin | Random | Accuracy | 37.3 % ± 0.5 % | 41.2 % ± 0.5 % | 55.0 % ± 0.5 % | 81.6 % ± 0.5 % |
| Xin | TF | Accuracy | 50.1 % ± 0.5 % | 63.3 % ± 0.4 % | 69.2 % ± 0.4 % | 85.5 % ± 0.4 % |
| Xin | Random | ARI | 0.102 ± 0.003 | 0.134 ± 0.003 | 0.240 ± 0.005 | 0.650 ± 0.007 |
| Xin | TF | ARI | 0.195 ± 0.004 | 0.333 ± 0.005 | 0.414 ± 0.006 | 0.706 ± 0.006 |
| Segerstolpe | Random | Accuracy | 40.3 % ± 0.7 % | 47.7 % ± 0.7 % | 58.0 % ± 0.7 % | 86.5 % ± 0.5 % |
| Segerstolpe | TF | Accuracy | 75.8 % ± 0.7 % | 80.2 % ± 0.5 % | 81.6 % ± 0.6 % | 90.7 % ± 0.4 % |
| Segerstolpe | Random | ARI | 0.174 ± 0.005 | 0.194 ± 0.005 | 0.303 ± 0.007 | 0.722 ± 0.008 |
| Segerstolpe | TF | ARI | 0.557 ± 0.009 | 0.610 ± 0.008 | 0.631 ± 0.009 | 0.798 ± 0.007 |
| Wang | Random | Accuracy | 38.0 % ± 0.5 % | 40.5 % ± 0.5 % | 53.2 % ± 0.5 % | 75.8 % ± 0.5 % |
| Wang | TF | Accuracy | 54.5 % ± 0.6 % | 65.6 % ± 0.5 % | 64.8 % ± 0.5 % | 78.6 % ± 0.5 % |
| Wang | Random | ARI | 0.113 ± 0.003 | 0.160 ± 0.004 | 0.266 ± 0.005 | 0.567 ± 0.007 |
| Wang | TF | ARI | 0.289 ± 0.006 | 0.396 ± 0.008 | 0.376 ± 0.006 | 0.599 ± 0.007 |

